# The Arch from the Stones: Understanding Protein Folding Energy Landscapes via Bio-inspired Collective Variables

**DOI:** 10.1101/2025.05.28.656575

**Authors:** Valerio Rizzi, Margaux Héritier, Nicola Piasentin, Simone Aureli, Francesco Luigi Gervasio

## Abstract

Protein folding remains a formidable challenge despite significant advances, particularly in sequence-to-structure prediction. Accurately capturing thermodynamics and intermediates via simulations demands overcoming timescale limitations, making effective collective variable (CV) design for enhanced sampling crucial. Here, we introduce a strategy to automatically construct complementary, bio-inspired CVs. These uniquely capture local hydrogen bonding—explicitly distinguishing protein-protein from protein-water interactions—and side-chain packing, taking into account both native and non-native contacts to enhance state resolution. Using these CVs in combination with advanced enhanced sampling methods, we simulate the folding of Chignolin and TRP-cage, validating our approach against extensive unbiased simulations. Our results accurately resolve complex free energy landscapes, reveal critical intermediates such as the dry molten globule and demonstrate agreement with reference data. This interpretable and portable strategy underscores the critical role of microscopic details in protein folding, opening up a promising avenue for studying larger, more complex biomolecular systems.

## Introduction

*Marco Polo describes a bridge, stone by stone. - But which stone supports the bridge? - asks Kublai Khan. - The bridge is not supported by this or that stone, answers Marco, but by the arch that they form. Kublai Khan keeps silent, thinking. Then he adds: - Why are you talking about the stones then? I am only interested in the arch. Polo replies: - Without stones there is no arch* [1]. The bridge described by Marco Polo to Kublai Khan in Italo Calvino’s Invisible Cities bears a remarkable resemblance to a folded protein with its global shape emerging naturally when all its interactions – the bridge’s stones – are in one place and no other.

Protein folding is a fundamental problem in biochemistry, with the structure-function paradigm being a cornerstone of biology and drug discovery [2–7]. Many decades have passed since the discovery that amino acid sequences determine how proteins fold into three-dimensional shapes [8], and significant progress has been made in the field until the recent recognition of the 2024 Nobel Prize in Chemistry awarded to the developers of Alpha Fold [9]. While the advent of Alpha Fold has made a significant leap forward in solving the problem of associating a structure to a sequence [9], several crucial aspects of protein folding are still largely unsolved and subject to intense research efforts [10], namely the microscopic details of how a protein folds [11, 12], the role of folding intermediates [13–16] and the systematic determination of thermodynamic and kinetic folding properties [17–20].

If all protein configurations were equally accessible and the folding probability landscape was flat like a golf course, the famous Levinthal paradox would apply [21, 22], making the folding time unfathomably long. On the contrary, at room temperature, energetic considerations discourage visiting an exceedingly large number of configurations, such as those where hydrogen bonds are unsatisfied [6, 7, 23]. The folding probability landscape must resemble a rugged funnel that is largest in the entropic unfolded basin and becomes narrower and narrower as one approaches the folded state [24–27].The metastable intermediates that populate the folding landscape can differ from each other by the smallest of details, such as the orientation of a side chain [23, 28] or the presence/absence of a single water molecule in a key position [29, 30]. Progress on this landscape is made by incremental stochastic steps that bring one closer and closer to the global free energy minimum, while crossing a myriad of small kinetic barriers [6].

Computer simulations are a powerful tool to shed light on the protein folding problem. In this respect, simplified models such as Gō models have greatly contributed to clarifying the general features of the protein folding landscape [31]. However, by constructions, these models forgo a detailed atomistic description. Atomistic molecular dynamics (MD) simulations would be well suited to the task, but are limited by the timescale problem [32]. Protein folding events tend to occur over hundreds of microseconds and, notwithstanding the continuous progress of computer hardware, reaching such long time scales remains a demanding task even today. Running long MD simulations on Anton [33, 34], a specialised computer for MD, produced groundbreaking results, generating folding trajectories of small proteins and predicting kinetic and thermodynamic properties close to the experimental ones [17, 35–38]. However, the inaccessibility of Anton to the wider academic community and the cost of designing specialised hardware limits the practical use of such approach.

Enhanced sampling algorithms provide a valid alternative to running extremely long MD on specialized hardware. In this respect, methods based on replica exchange techniques [39–43] and on collective variables (CV) have been used on protein folding with some success [20, 44, 45]. However, even collective variable based enhanced sampling methods such as Metadynamics [46, 47] and On-the-fly Probability Enhanced Sampling (OPES) [48–50] are limited by the quality of the chosen CVs that need to properly encode the phenomenon of interest [51, 52]. Recently, we have developed a hybrid method called OneOPES [53] that, by exploiting the power of OPES Explore [50] and of agnostic techniques such as OPES MultiThermal [49] and replica exchange [54], makes the requirement to develop optimal CVs a less severe but still very valuable endeavour.

Good quality CVs must first be able to distinguish the end states - the unfolded and folded state - then to capture possible intermediates along the folding path and to encode as many as possible slow degrees of freedom (DOFs) that yield the folding/unfolding kinetic bottlenecks. A number of folding CVs have been developed over the years, from the ones that capture global properties like the root mean square deviation (RMSD) away from a state, the radius of gyration [44] or the number of turns of alpha helices [55] to CVs that combine a number of local features such as interatomic distances or dihedrals according to different criteria [45, 56–64].

One of the main limitations shared by existing folding CVs is degeneracy: it is extremely difficult for a CV to resolve well the multitude of metastable states relevant to folding, especially in close proximity to the native folded state. Accelerating degenerate CVs slows down recrossings, leads to hystheresis and hampers convergence. CVs that encode global properties are intrinsically affected by this problem as they typically cannot resolve the microscopic interactions that characterise the folded basin. For example, CVs that exclusively focus on the protein’s backbone such as interatomic distances between C*α*s or dihedrals do not resolve the reciprocal position of the side chains and do not have an explicit focus on capturing hydrogen bonds. The degeneracy problem increases in severity with system size: attempts to use CV-based enhanced sampling to converge the folding free energy on proteins larger than the standard 10 residue Chignolin mini-protein [65, 66], such as the 20-residue TRP-cage mini-protein [67], are few [68] and less successful [59].

In this context, the elephant in the room may be the role of the solvent molecules that account for the vast majority of the atoms in a simulation box. Many interactions play a role in the folding of globular proteins, among which electrostatic interactions and steric hindrance, but hydrophobicity has emerged possibly as the leading factor [69]. The egress of water molecules from the protein hydrophobic core and the replacement of solvent-mediated hydrogen bonds with direct protein-protein hydrogen bonds are key steps where water plays an essential role. Accelerating the dynamics of water molecules at relevant positions is an idea that is gaining traction [70–72] and we have recently proposed specialised water CVs in different scenarios, such as ligand binding [73–77], ion transport [78], and local conformational changes [79, 80].

In this work, we develop a bottom-up strategy that builds two complementary CVs, starting from the selection of bio-inspired features. One CV focuses on capturing individual high-energy microscopic interactions - hydrogen bonds - that identify and distinguish protein-protein from protein-water bonds, while also encoding angular information. The other captures mesoscale contacts between side-chains, encoding compactness. The features are automatically collected from short unbiased end-state trajectories and filtered using a Linear Discriminant Analysis (LDA)-like criterion [81]. For both CVs, a sharp distinction between the true folded state and defective metastable states is achieved by summing native contacts and explicitly subtracting non-native contacts. Finally, the two CVs are intuitively merged as a simple linear combination of the features with unitary coefficients.

We use such CVs to investigate the folding free energy of two well-known miniproteins from the seminal paper of Lindorff-Larsen et al. [17]: Chignolin [65, 66] and TRP-cage [67]. The references for all simulations are self-generated long unbiased trajectories of 300 *µ*s for Chignolin and 200 *µ*s for TRP-cage, which are made available for future reference and CV development. The enhanced sampling simulations are run in quintuplicate to ensure reproducible results. For Chignolin, both single replica OPES and OneOPES simulations converge to analogous results, with OneOPES being faster and more robust. For TRP-cage, OneOPES calculations are in good agreement with the reference. We analyse the resulting folding landscape, focusing on the metastable intermediates states leading to the native state, such as the elusive dry molten globule (DMG) [13, 14, 16, 82].

The convergence speed and quality across the set of examples, combined with the intuitive and portable nature of the CVs, support our claim that the construction of highly optimised microscopic features represents the crucial element to be accelerated in folding simulations, confirming the paradigm that folded structures emerge naturally from the fulfilment of microscopic contacts. Our automated strategy is well suited for combination with modern machine learning strategies [83–85] and can pave the way for in-depth studies of larger and more complex proteins.

## Results

### A bottom-up strategy for folding featurisation

An effective approach to designing CVs involves initially observing and defining a number of features of the system under investigation, and then combining these features into CVs at a later stage. To accelerate the process of interest in enhanced sampling, CVs and features must first be able to distinguish states and follow the process through phase space. Ideally, they should align themselves with the lowest free energy path and approximate the committor [62, 63].

While a rather large number of approaches for combining features exists, the features’ search and optimization is often neglected, despite being a more fundamental aspect than the combination stage. In fact, an incomplete or suboptimal choice of features that ignores crucial DOFs has no chance of producing an optimal CV, even when coalesced with the most sophisticated combination techniques.

A common choice of features is the set of all interatomic distances or contacts between heavy atoms [45, 64, 86–89]. This is often an excellent option, especially in low-density gas phase systems, as it captures a large amount of information and can distinguish well microstates while respecting translational, rotational, and permutational symmetries. However, in protein folding, this choice can be suboptimal. For instance, the number of distances grows quadratically with the number of atoms, requiring a preliminary selection of a subset of atoms for manageability. The distances between C*α*s are a common choice, but they do not encode any information about the side-chains’ movements and do not explicitly detect native and non-native contacts (NNCs). Such information is essential for resolving the intermediate metastable states, or kinetic traps, that populate the rugged folding landscape and requires a level of microscopic detail that C*α* distances alone cannot furnish.

Finally, and perhaps most importantly, any set of protein-protein distances neglects the role of the solvent. This, frequently considered less important, holds a significance at least equivalent to that of the protein atoms, especially in satisfying hydrogen bonding. In fact, the energy contribution of a *single* hydrogen bond lies in the range of a few kcal/mol, which is comparable to the whole folding free energy difference of many small proteins. Consequently, the total number of hydrogen bonds tends to be a highly conserved quantity throughout a folding/unfolding process [6, 23, 90]. Breaking an intra-protein hydrogen bond simply by stretching the corresponding protein-protein distance risks crossing high-energy, improbable states, as no alternative hydrogen bond is promptly available for replacement. Instead, simultaneously driving a water molecule into the correct position to replace the cleaved hydrogen bond helps in reducing the free energy cost of crossing the barrier, thereby better approximating the minimum free energy path.

Because of these considerations, in this paper we propose a bottom-up strategy to build folding CVs starting from picking and optimising two key sets of features:

- the formation and cleavage of H-bonds 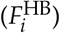;
- the relative packing of the side chains 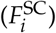.

Both are crucial to ensure a comprehensive description of secondary structures (*α* helices and *β* sheets) and disordered portions. In the final CVs, features that capture native contacts that exist in the folded basin 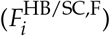 are summed with coefficient 1, while those that capture contacts from the unfolded basin 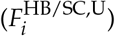 have coefficient −1. The two resulting CVs are simply

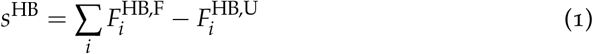

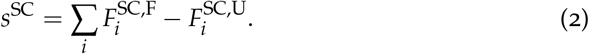

Individual features 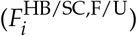 act as a switch, going from ≈ 1 when a native contact is present to ≈ 0 when no contact is present until ≈ −1 in case of a nonnative contact. In the Methods section, we detail the construction of both sets, their filtering process, and their combination into CVs. Our workflow requires only two short unbiased trajectories, one in the folded basin and one in the unfolded basin, and is fully automated and available as a Python script. The script output consists of PLUMED [91] files, ready for either trajectory post-processing or enhanced sampling simulations. Additional details can be found in the Supporting Information (SI).

### Chignolin

At first we simulate the folding of a well-known system that has been widely used in method testing over the years, the double mutant CLN025 of the Chignolin miniprotein [65, 66]. Our simulation input parameters match when possible the ones from Ref. [17] (see the Methods section for more details). We use as a reference a self-generated set of 3 unbiased trajectories, each 100 *µ*s long (see Fig. S3) to calculate our reference folding free energy Δ*F* = −1.66*±*0.16 kcal mol^−1^. We observe a slight shift compared to the reported value of −0.9 kcal mol^−1^ [17]. We believe this to be due to subtle differences in the implementation of the force field in different MD codes.

To investigate where the energy shift emerges, we evaluate the enthalpy difference Δ*H* by partitioning the total energy of our unbiased NVT trajectories into folded and unfolded basins (see Fig. S4). Our resulting Δ*H* = 7.1*±*0.3 kcal mol^−1^ is shifted with respect to the value reported in Ref. [17] Δ*H* = 6.1*±*0.1 kcal mol^−1^ by a similar amount as the folding free energy Δ*F*, giving credit to our hypothesis of slight force field discrepancies.

To test our CVs in the context of enhanced sampling, we first perform single replica simulations of the system biasing *s*^HB^ and *s*^SC^ with OPES [48] (see Fig. 2 and Fig. S5, 7). This is already a rather stringent CV test where suboptimal CVs are known to display hysteresis and evident discrepancies in the estimated Δ*F* between independent simulations. Instead, our Δ*F*(*t*) estimate is consistently aligned in all replicas with the expected value after about 400 ns (see Fig. 2 (a)) and remains rather constant throughout the rest of the simulations, giving a final estimate of Δ*F* = −1.81*±*0.20 kcal mol^−1^. The FES profile along the commonly used RMSD over the C*α* atoms is in good agreement with the unbiased reference (see Fig. 2 (b)).

We then perform analogous OneOPES simulations and observe a faster convergence and an even stronger agreement with the reference (see Fig. 2 (c,d) and Fig. S6, 7), as well as a more consistent behaviour among the independent simulations than in the single replica case. This is expected as the use of explicit replica exchange and the bias over additional CVs helps to accelerate degrees of freedom that are not directly sped-up by the main CVs. Our final folding free energy estimate is Δ*F* = −1.64 *±* 0.15 kcal mol^−1^ and compares well with the reference (see Tab. 1).

**Table 1.** Free energies of folding Δ*F* of the reference unbiased systems and the OneOPES simulations.

| | $\Delta F$ (kcal mol<sup>-1</sup>) | |
| --- | --- | --- |
|  | OneOPES | Reference |
| **Chignolin** | $-1.64 \pm 0.15$ | $-1.66 \pm 0.16$ |
| **TRP-cage** | $0.91 \pm 0.16$ | $0.86 \pm 0.27$ |

### TRP-Cage

We now apply our method to study TRP-cage, a more complex mini-protein that has been the focus of a number of experimental [39, 67, 92, 93] and computational studies [20, 35, 38, 56, 59, 68, 94]. We study the TC10b K8A single-point mutant that features in the seminal work by Shaw et al. [35] and we setup the system to match the one from Ref. [38] (see also the Methods section).

TRP-cage contains 20 residues and features three distinct secondary structure elements: an *α*-helix spanning residues 1–9, a 3_10_ helix from residue 11 to residue 13, and a polyproline II helix between residues 17–19 (see Fig 1 (d)). These elements assemble into a compact tertiary structure that encapsulates a central tryptophan residue (W6), forming a primitive hydrophobic core in conjunction with residues Y3, G11, P18, and P19. Its richer topology compared to Chignolin makes TRP-cage a challenging and informative benchmark for folding simulations. Furthermore, the folding mechanism of TRP-cage can be regarded as a surrogate for the folding of larger proteins, as the rise of its well-defined hydrophobic core entails a hydrophobic collapse and the formation of molten globule intermediate states [95]. We use an extensive self-generated unbiased simulation of 200 *µ*s as a reference, from which we estimate the folding free energy to be Δ*F* = 0.86 *±* 0.27 kcal mol^−1^ (see also Fig. S8). After building the *s*^HB^ and *s*^SC^ CVs, we run five independent OneOPES simulations (see Methods and Fig. S9, S10 for details) and we compare their outcome with the unbiased reference. The folding free energy from the OneOPES simulations, in Fig. 3(a), displays a remarkable agreement with the reference from 500 ns onwards. The resulting FES projected on the RMSD over the C*α* atoms, in Fig. 3(b), matches the reference well, with the error peaking in the transition state region, as seen previously with OneOPES applied on other systems [53].

**Fig. 1.**
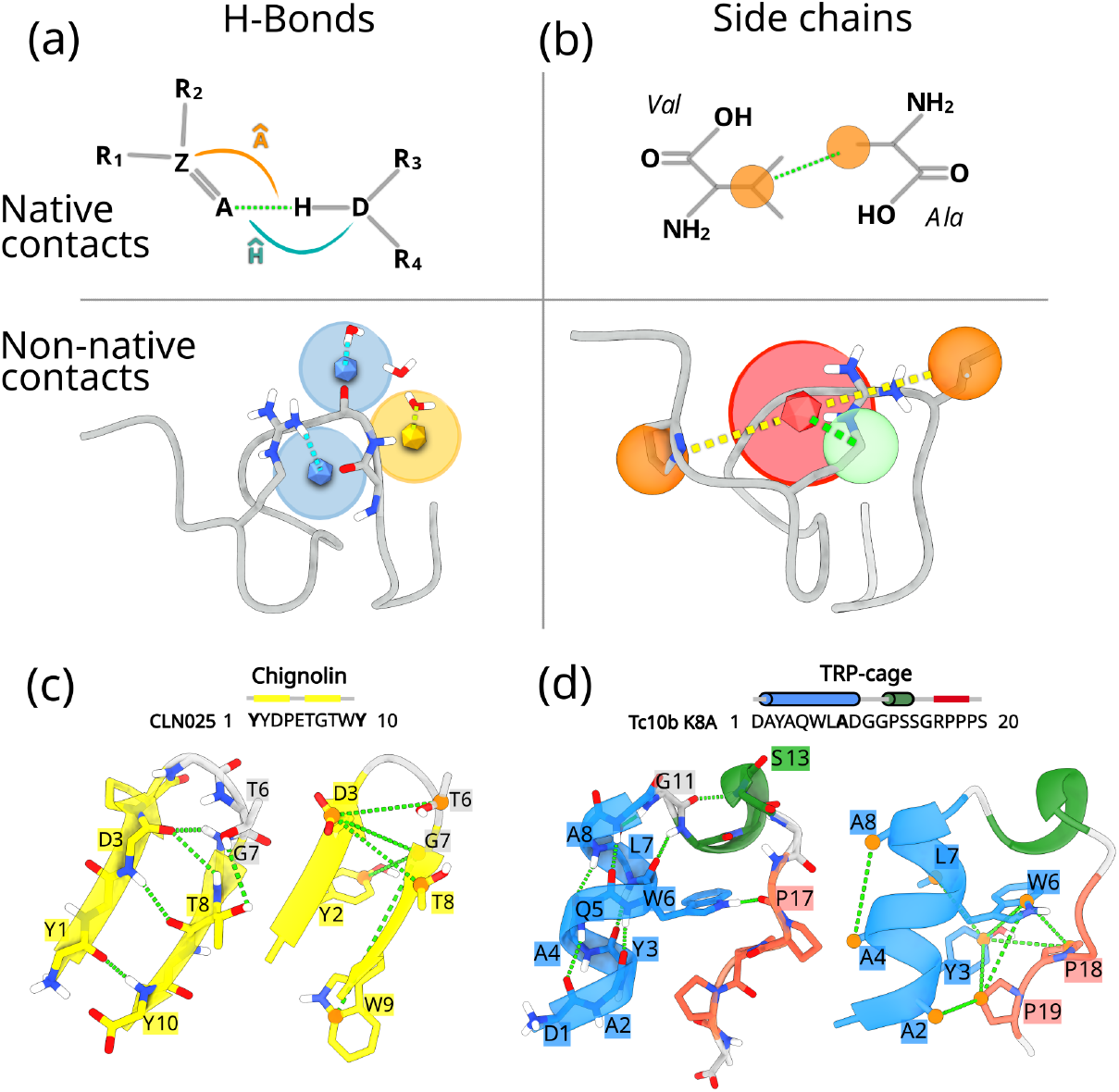
Feature design and systems studied. **(a)** Illustrative definition of an H-Bond feature considering the bond angular components for native contacts and the protein-protein and protein-water interactions for non-native contacts. **(b)** Illustrative definition of the side chain feature with both a native and a nonnative interaction between centres of mass of side chains. **(c**,**d)** Sequence and 3D native structures of Chignolin and TRP-cage, respectively. *α*-helices are represented in blue, 3_10_-helices in green, *β*-helices in yellow, and random coils in grey. The green dashed lines highlight the most relevant H-bonds contacts (left) and the most relevant side chain contacts (right).

**Fig. 2.**
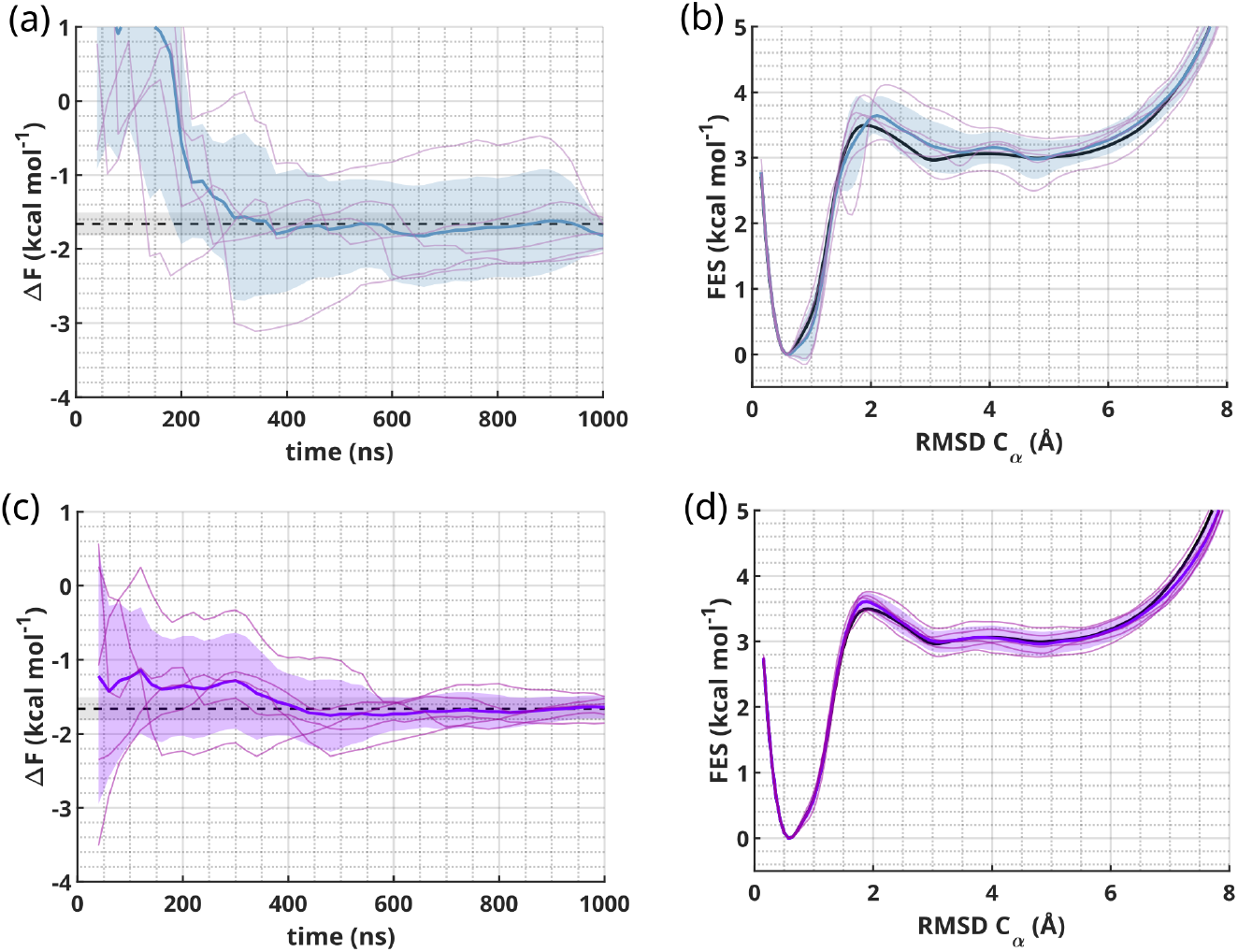
Chignolin folding results. **(a**,**c)** Chignolin free-energy difference Δ*F* between the folded and unfolded states over time from five independent OPES **(a)** and OneOPES **(c)** trajectories. The reference value is displayed as a black dashed line, the trajectories’ average value in solid color, blue and purple, respectively, and the standard deviation in semi-transparency. **(b-d)** 1D FES as a function of the RMSD over the C*α* atoms for the OPES **(c)** and the OneOPES **(d)** simulations. In solid black line we show the reference FES, in solid colour, blue and purple, respectively, we report the average of the five replicas, and in semi-transparency their standard deviation.

**Fig. 3.**
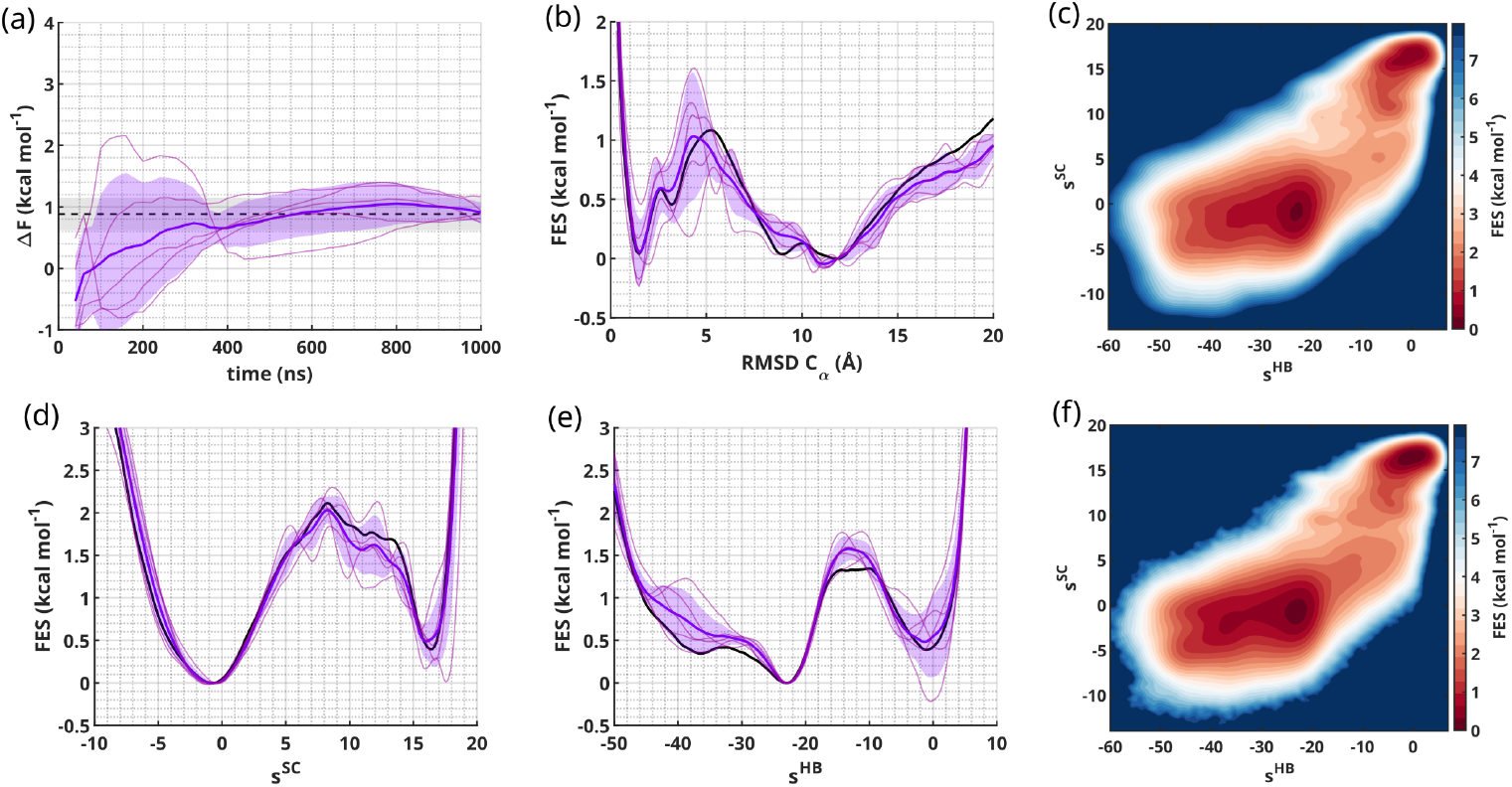
TRP-cage folding results. **(a)** TRP-cage free-energy difference between the folded and unfolded states over time from five independent OneOPES trajectories. The reference value is displayed as a black dashed line, the trajectories’ average value in solid purple and the standard deviation in semi-transparency. **(b)** 1D FES as a function of the RMSD over the C*α* atoms with the reference in solid black, the OneOPES average in solid purple and the standard deviation in semi-transparency. **(c**,**f)** 2D FES over *s*^HB^ and *s*^SC^, with the average value over the OneOPES independent trajectories in **(c)** and the reference in **(f). (d**,**e)** 1D FES as a function of *s*^SC^ **(d)** and *s*^HB^ **(e)**, with an analogous colour scheme as the one in panel **(b)**.

The 2-dimensional FES projected on the *s*^HB^ and *s*^SC^ CVs, in Fig. 3(c), closely matches the one built from the reference in Fig. 3(f) and reveals important topological features that deserve a more in-depth analysis. The two main minima corresponding to the unfolded (*s*^HB^ ≈ −25, *s*^SC^ ≈ −2) and the folded basins (*s*^HB^ ≈ 0, *s*^SC^ ≈ 18) are surrounded by plateau-like regions that contain a number of other metastable states. The reweighting of the 2D FES along its 1-dimensional components, reported in Fig. 3(d,e), reveals that, while the projection over *s*^SC^ is dominated by the two main minima, the one over *s*^HB^ shows an unfolded basin made up of two components.

To probe the source of such bifurcation and further investigate the nature of metastable states, we extract configurations belonging to different regions of the 2D FES and analyse their secondary structure content. As shown in Fig. 4, we single out six population ensembles, respectively located in:

- the unfolded basin, state U and M (standing for unfolded and misfolded states);
- the transition state region, state WMG (standing for wet molten globule);
- the folded basin, states F, F2 and DMG (standing for folded and dry molten globule states).

**Fig. 4.**
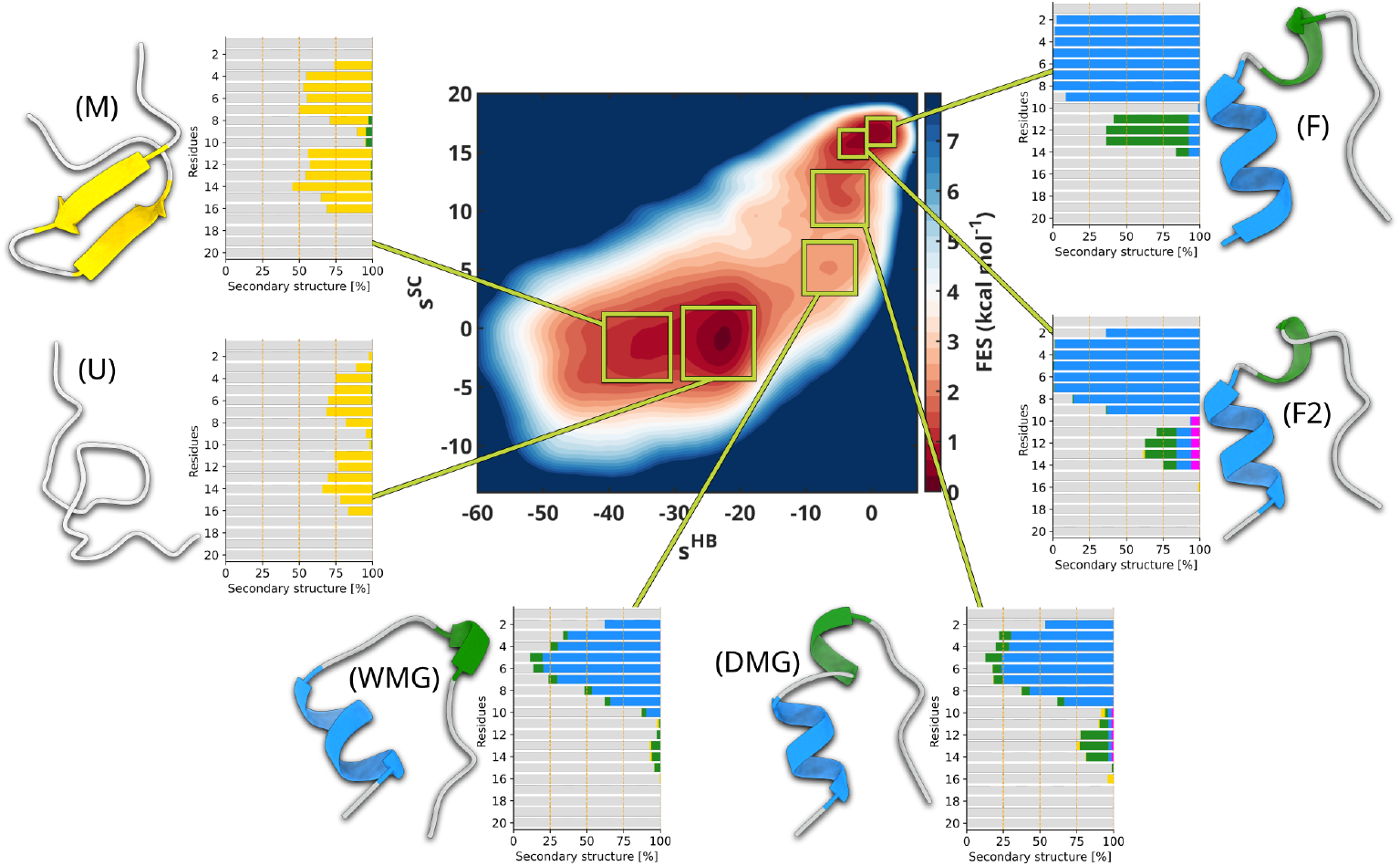
Trp-Cage folding landscape analysis. Six conformational ensembles (M, U, WMG, DMG, F2 and F) extracted from the corresponding regions of the OneOPES TRP-cage 2D FES highlighted by green squares. For each ensemble, on the left we present a representative 3D structure and on the right a histogram summarising the secondary structure frequency for each amino acid. *α*-helices are coloured in blue, 3_10_-helices are coloured in green, *π*-helices in magenta, *β*-helices in yellow, and random coils in grey.

State U is located in the region −30 ≲ *s*^HB^ ≲ −20, −4 ≲ *s*^SC^ ≲ 2 and corresponds to a fully unfolded ensemble of states lacking stable secondary structure, though retaining a modest degree of local order in the regions where the *α*-helices could form. In contrast, and perhaps surprisingly, state M, located in the interval −40 ≲ *s*^HB^ ≲ −30 and in a similar *s*^SC^ range as the above states U, presents a misfolded and partially folded motif, a *β*-turn-*β* conformation. To further assess the differences between U and M, we analyse the per-residue solvent-accessible surface area (SASA) and contact frequencies (see Fig. S11). While the SASA values for the two states are comparable, state M clearly displays contact patterns characteristic of *β*-hairpins, with increased interactions between distal residues.

The capability of *s*^HB^ to distinguish U from M is not trivial and is intrinsic to its construction where NNCs are subtracted from the native contacts. While disordered structures contribute almost nothing to *s*^HB^, misfolded ones contribute negatively, and the more structured they are, the more negative is their contribution. The capacity to place misfolded structures on one end of the CV value range is a key quality for a good CV to be used in enhanced sampling as it contributes to clean up the intermediate region from dead ends and kinetic traps. If one looks at a standard CV such as the RMSD over the C*α* atoms (see Fig. S10), its value in state M is indeed mixed with the value of intermediate states that appear in the transition state region. When biased, such CVs might manage to lead the system from U to the transition region, but would probably be trapped in M for a long time. Further progress towards state F would be hampered by the necessity to first go backwards to the disordered state U before being able to move forward to F. The pitfalls of using such problematic CVs in the context of a simple but pathological two-dimensional potential are well illustrated by Invernizzi *et al*. in Ref. [50].

In the intermediate region of the free-energy surface −12 ≲ *s*^HB^ ≲ −2, 4 ≲ *s*^SC^ ≲ 8, we identify a conformational ensemble that we associate with a wet molten globule (WMG). Structural analysis reveals the emergence of an *α*-helix spanning residues 3–8, supported by increased contact frequencies within this segment. Notably, the dominant interactions involve Y3, W6, and L7, likely marking the early nucleation site of the hydrophobic core (see Fig. S11). Despite this local ordering, the per-residue SASA remains high and comparable to that of U and M, indicating that WMG is still fully solvated.

In the folded region, we identify three states F (folded), F2 (misfolded) and DMG (dry molten globule) that are nearly indistinguishable when characterized using coarse collective variables such as RMSD or AlphaRMSD (see Fig. S10). On the contrary, the atomistic detail encoded in *s*^HB^ and *s*^SC^ can resolve the subtle structural differences among them. While states F and F2 feature a well-formed *α*-helix spanning residues 2–9, state DMG presents a slightly less ordered helical region that resembles the one present in WMG. The main difference between WMG and DMG lies in per-residue SASA values that significantly drop, especially for W6, G11, and P19, i.e. the key residues involved in hydrophobic core formation. Their decreasing solvent exposure indicates water expulsion and core consolidation, which ultimately lead to the formation and the stabilisation of the native-like fold. The SASA values further decrease from DMG to F2, reflecting an increasing compactness.

F and F2 closely resemble each other when looking at SASA values but differ locally in secondary structure. Only state F exhibits persistent folding of the helix’s N- and C-terminal ends, as it is expected in a native folded structure. In the region encompassing residues 10–14, F2 shows diverse helical content, including transient 3_10_ and *π*-helix features, whereas F keeps a stable 3_10_ helix. In Fig. S11, we repeat the same analysis over the same metastable states on the reference unbiased trajectory. The secondary structure content of each basin is largely in agreement with the one that we find in the OneOPES trajectory.

Our results on TRP-cage highlight the power of the *s*^HB^ and *s*^SC^ CVs to deliver both accurate thermodynamic estimates of the folding free energy and high-resolution structural information. By capturing subtle structural and hydration differences, our approach is able to accelerate the relevant slow degrees of freedom and offers valuable insights on the ensemble of states along the folding path.

## Discussion

In this work, we introduce a bottom-up strategy to design physically interpretable CVs capable of grasping the intricate details of protein folding energy landscapes, both at the microscopic and mesoscale level.

A key strength of our approach lies in the complementary nature of the two collective variables. *s*^HB^ provides a comprehensive atomistic description of hydrogen bonding, capturing secondary structures at a high-resolution and efficiently promoting their formation and dissolution in enhanced sampling. Conversely, *s*^SC^ captures side-chain packing at a coarser scale level, effectively describing and driving hydrophobic collapse and tertiary structure formation. Together, these CVs bridge the gap between microscopic and mesoscale, enabling enhanced sampling simulation to reversibly sample the conformational space until convergence. This synergy proves to be particularly powerful in detecting intermediate and near-native states that might otherwise be overlooked using traditional CVs alone.

We integrate these CVs in the OneOPES enhanced sampling framework and investigate the folding mechanisms of two mini-proteins: Chignolin and TRP-cage. Our approach yields free energy landscapes in strong agreement with extensive unbiased simulations. These results underscore the robustness and accuracy of our CVs in quantifying folding thermodynamics. In the case of TRP-cage, we dissect the structural features of relevant basins, resolving and characterising unfolded, intermediate, and folded states. Our analysis reveals distinct folding routes and highlights the role of hydrophobic core formation in native structure stabilization.

The success of our automated strategy in generating these high-resolution CVs from short unbiased end-state trajectories is promising. While previous CV-based enhanced sampling studies have often struggled with proteins larger than Chignolin, our results with TRP-cage demonstrate the potential of our approach for more challenging systems. To support reproducibility and foster further CV improvements, we make available the full trajectories of the reference unbiased MD simulations that we generated on both systems. We wish that these datasets may serve as benchmark references for existing strategies and future developments. The current validation on two mini-proteins paves the way to study larger and more complex protein systems with different folding mechanisms or structural characteristics. Assessing the generalisability of our feature set on such systems will be crucial to demonstrate the scalability of the approach.

Looking ahead, this work opens several exciting avenues. The interpretable and portable nature of our CVs makes them highly suitable for integration with modern machine learning techniques to further enhance CV optimization. Applying this methodology to larger proteins will be a key next step to explore phenomena like domain folding and local conformational changes. The detailed free energy landscapes generated would provide deeper insights into folding pathways, the nature of transition states, and the thermodynamics and kinetics of misfolding and aggregation processes implicated in disease.

Furthermore, the principles underlying our CV construction strategy – particularly the emphasis on distinguishing native versus non-native interactions and the explicit role of solvent – could be adapted to study more complex biomolecular processes, such as protein-ligand interactions, RNA folding, and other large-scale conformational transitions. This approach thus offers a valuable and extensible framework for advancing our understanding of the molecular mechanisms governing biological systems.

## Methods

### H-bond feature definition

An H-bond is an electrostatic interaction between a hydrogen atom *H* that is covalently bound to a more electronegative *Donor* atom *D* and an H-bond *Acceptor* atom *A* that must be electronegative and must possess a lone pair of electrons, covalently bound to atom *Z* (see Fig. 1 (a)). In this section, we focus on a single H-bond between atoms *H*_*i*_ and *A*_*j*_ extracted from an unbiased trajectory of one of the end states (indexes are omitted for simplicity in Fig. 1 (a) and in the text below) and go through the corresponding H-bond feature 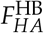 construction. The selection and filtering process of the H-bonds is discussed below.

Hydrogen bonding arises from the overlap between *H*’s orbital and the sp^3^ hybrid orbitals of *A*, which exhibit spatial anisotropy. While the strength of an H-bond is mainly modulated by the 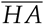 distance, it also depends on the orientation of its constituent atoms. We monitor the former through a coordination function *C*_*HA*_ whose value ranges from 1 when the bond is present to 0 at longer distances. We use a long-range switching function (SWF) of the following functional form

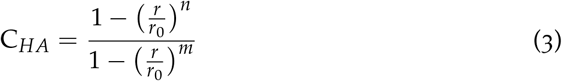

where 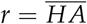, *r*_0_ = 3 Å, *n* = 6 and *m* = 8 (see Fig. S1, in blue).

The atomic orientation is encoded by two functions Γ_H/A_ (see Fig. 1a) which are equal to 1 when the angles 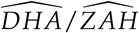, centred on atoms *H* and *A* respectively, are flat and decrease as the angles bend. These functions are simply defined as a ratio of distances

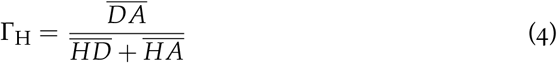

and

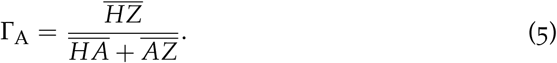

We define variable

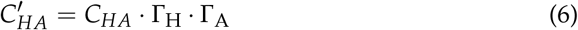

as an indicator of H-bond strength that takes into account both bond length and angular contributions (see Fig. S2 for illustrative plots of 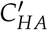 as a function of bond length and angle).

In an ideal *α*-helix or *α*-sheet backbone, native hydrogen bonds align perfectly, maximizing their strength. However, thermal fluctuations, solvent exposure and environmental interactions introduce deviations, weakening the bonds and increasing their flexibility. 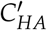 is a powerful tool for classifying H-bonds according to their length and orientation. If its average value in the examined unbiased trajectory is above an arbitrary threshold of 0.6, we consider the H-bond to be strong and name it *hard*. Otherwise, we name a wobblier H-bond *soft*. For hard H-bonds, we bring forward to the H-bond feature definition the full angular information contained in 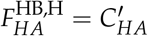, while for soft H-bonds we revert to a simpler form 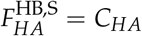.

We further classify an HB feature in terms of its specialisation in capturing a bond in either the folded or the unfolded state. To do so, we calculate the mean value of the H-bond feature 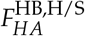 during the folded and unfolded unbiased trajectory. If the feature has a higher mean value in the folded state, it captures a native H-bond and we call it 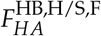, otherwise we call it 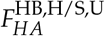. Naturally, as the folded state is more structured, the number of 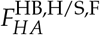 features tends to be larger than the number of 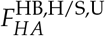.

For an H-bond feature to work efficiently in driving folding and unfolding events, it must contain information not only about a native bond being present, but also about other competing non-native bonds being absent. As a native H-bond contact of a protein cleaves during protein unfolding, energetic considerations dictate that it must be instantly replaced by a non-native interaction with other protein H-bond donors/acceptors or with water molecules. This bond replacement mechanism can be intuitively included into a feature by using opposite signs [96] for the two categories of native and non-native bonds.

We evaluate non-native interactions with protein atoms only for the folded-state-focused features 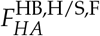 with the aim to boost the capability of the resulting H-bond CV to strongly distinguish the folded state from the variety of semi-folded metastable states that resemble it. Such a need is not present in the unfolded basin that naturally includes a number of very different structures. Sharply determining the formation of a NNC and filtering it from the surrounding noisy environment is crucial. The formation of NNCs can take place on both ends of the HB, namely either around atom *H* or *A*. To take both into account, we first dynamically determine two virtual atom positions along the 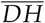 and the 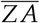 vectors, 2.5 Å away from *D* and *Z* (see Fig. 1(a)). We call these virtual atoms *V*_*H*_ and *V*_*A*_, respectively.

To evaluate the coordination number *C*_*i*_ between *V*_*H*_ and all possible non native protein acceptor atoms and between *V*_*A*_ and all possible non native protein donor atoms, we use the following short-ranged SWF

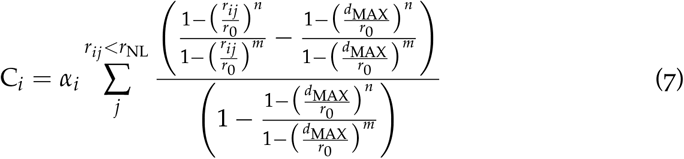

where *α*_*i*_ is a normalization factor and *r*_*ij*_ represents the distance between a virtual atom *i*, either *V*_*H*_ or *V*_*A*_, and all non-native protein acceptor/donor atoms *j*. This SWF goes smoothly to zero at distance *r*_*ij*_ = *d*_MAX_ and is null in the buffer zone *d*_MAX_ *< r*_*ij*_ *< r*_NL_ where *r*_NL_ is the neighbour list radius. We set *r*_0_ = 3.5 Å, *n* = 2, *m* = 10, *d*_MAX_ = 5 Å, *r*_NL_ = 8 Å (see Fig. S1, in red), and update the neighbour list every 20 steps. The short-rangeness of the SWF is essential in picking up the signal of specific NNCs arising in close vicinity to *i* and in reducing the noise from atoms further apart.

In the case of non-native interactions with water, we calculate them for both folded-state-focused and unfolded-state-focused features, but only for hard H-bonds *F*^HB,H,F^ that present a well defined position where a water molecule would stay. We evaluate the coordination number between *V*_*H*_ and all water oxygen atoms and between *V*_*A*_ and all water hydrogen atoms using the same SWF from Eq. 7. As before, we set *r*_0_ = 3.5 Å, *n* = 2, *m* = 10, *d*_MAX_ = 5 Å, *r*_NL_ = 8 Å(see Fig. S1, in red), and update the neighbour list every 20 steps.

Ideally, we would like the value of an HB feature to go from about 1 when the native contact is present to about − 1 when a NNC replaces it. To achieve so, the NNC coordination number must be normalised so that its value compensate and does not exceed the one of the native contact. In the case of a NNC calculated between *V*_*H*_ and water oxygen atoms 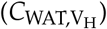 we use a normalisation factor *α*_*i*_ = 1/8, while in the case between *V*_*H*_ and protein acceptor atoms 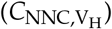 we use a normalisation factor of 1. On the other hand, for NNC calculated between *V*_*A*_ and the more numerous water hydrogen atoms 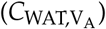 we use a normalisation factor of 1/16 and finally in the case between *V*_*A*_ and protein donor atoms 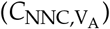 we use a normalisation factor of 1/2.

All in all, the final form of a folded-state-specialised H-bond feature is

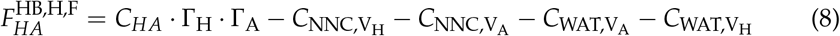

for a hard H-bond and

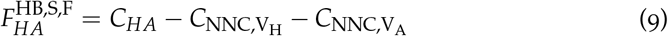

for a soft H-bond.

Instead, an unfolded-state-specialised feature is

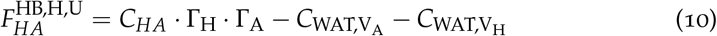

for a hard H-bond or simply for a soft H-bond.

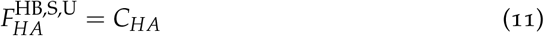

In summary, the difference between folded-state-specialised features 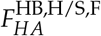 and unfolded-state-specialised features 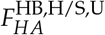 lies in the presence of the NNC terms with respect to protein acceptors/donors only in the folded-state-specialised case. These terms help to sharpen the resolution of the CV in the key region in close vicinity to the folded state, while the same would not be helpful in the more unstructured unfolded basin.

### Side chain packing feature definition

The second feature that we craft, *F*^SC^, specialises in capturing side chain packing through the relative orientation of the residues. It is complementary to the H-bond feature as it is aimed at distinguishing states that may share H-bonds contacts and secondary structure, but have a different degree of compactness. Its components describe individual native contacts between side chains and distinguish by construction spurious non-native contacts that characterise metastable states.

We describe the side chains at the mesoscale level with the distances of their centre of mass (see Fig. 1 (b)). In the special case of side chain-free glycine residues, the side chain COM is replaced by its C*α* atom. In analogy with our treatment of *F*^HB^, we focus on the case of one contact between hypothetical residues *X* and *Y* and leave the contact selection and filtering process to the next section.

First, we transform the 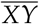 distance into a contact *C*_*XY*_ through a SWF of the form contained in Eq. 3. We use *r*_0_ = 8 Å, *n* = 4 and *m* = 8. This SWFs is rather long-ranged as it is focused on picking up a single specific contact between *X* and *Y*, even when they are far apart (see Fig. 1 (b) and the curve in green in Fig. S1). Its long-rangedness is fundamental to pull *X* and *Y* together when going from the unfolded basin to the folded basin in enhanced sampling. Depending on the mean value of *C*_*XY*_ in the unbiased trajectories, the resulting feature would be either folded-state specialised 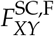 or unfolded-state specialised 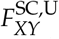.

To better distinguish metastable states from the end states and reduce overall degeneracy in the frustrated space of side chain packing [28], we also include information about NNC, with a negative sign in a similar fashion as we do for *F*^HB^. During the trajectories, we define a virtual atom *V*_*XY*_ at the geometrical centre between *X* and *Y*. We calculate the contact between *V*_*XY*_ and all other side chain COMs beside *X* and *Y* (see Fig. 1 (b)). To reduce noise, we use here the short-ranged SWF from Eq. 7 with *r*_0_ = 3.5 Å, *n* = 2, *m* = 10, *d*_MAX_ = 5 Å, *r*_NL_ = 8 Å, and a neighbour list update every 20 steps. We call the resulting contact 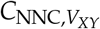.

The final form of the side chain packing feature is then

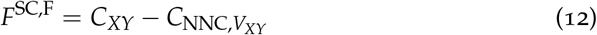

for a folded-state-specialised feature and

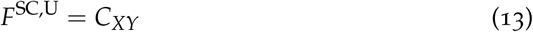

for an unfolded-state-specialised feature.

### Filtering features

Up to now, we have described in detail how to design features assuming that we already know which ones are relevant. Here we take a step back and show how to automatically select them. This selection process is crucial for reducing the very large space of possible features and picking out only the most relevant ones, thus filtering unwanted noise in the CVs to be generated.

The starting point is two short unbiased trajectories of the folded state F and the unfolded state U, indicatively 100 ns each totalling at least 1000 independent configurations.

Our leading criterion is to rank and filter the features according to their discriminative power between the two end states via an LDA-like score. Through successive steps, we apply a filtering strategy to progressively identify the most relevant H-bond and side-chain packing features and discard those that do not contain a sufficient level of information.

For the H-bond features, we first perform a rough preliminary selection by gathering all the possible combinations *ij* between any hydrogen atom *H*_*i*_ bound to an H-bond donor *D*_*i*_ and any H-bond acceptor *A*_*j*_ and calculating their distance. At first we keep only the combinations where a long-lived H-bond exists by retaining only the features in which the mean distance in either the folded/unfolded basin 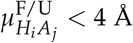. Out of these, we keep only those where 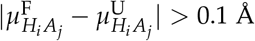, hence discarding all the H-bonds that are equally stable in both end states. These filtering steps significantly reduce the number of H-bonds feature candidates by discarding all contacts that do not generate an H-bond in either state or that generate one in both states.

To classify whether an H-bond is hard or soft, we evaluate its corresponding descriptors 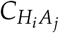 and 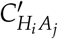 as defined in Eq. 3 and Eq. 6, respectively. If the average value of 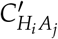 in either of the end states is above a threshold value of 0.6, then the H-bond is classified as hard and is retained. If not, we revert to the simpler contact 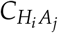. If its average value 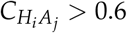 on either state, the H-bond is classified as soft and is also retained. If not, the H-bond is discarded.

Out of the remaining features, we evaluate their standard deviation over the end states 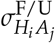 and calculate their discriminative power

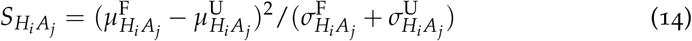

according to an LDA-like criterion. We keep features with a discriminative power 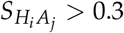.

Features are further classified according to the sign of 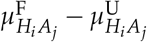, with a positive sign indicating features that are folded-specialised and, conversely, a negative sign for unfolded-specialised.

In the calculation of NNC that lead to protein-protein hydrogen bonds (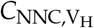 and 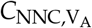), we include in the summation all possible donor-acceptor combinations, excluding the non native contacts that are in close vicinity to the given native one in the folded state i.e. with an average distance less than 4 Å.

For the side chain packing features, the filtering procedure is analogous and simpler. We take into account only non adjacent side chain contacts, excluding those between residues *X*_*i*_ and *Y*_*i*±1_. Out of these features, we keep all those that have a value 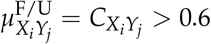 in either state and a discriminative power 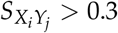.

### Additional hydration CVs

In the OneOPES strategy, additional CVs can be biased in higher order replicas in order to accelerate supplementary DOFs and boost convergence. In this context, given the importance of the protein solvation and the hydrophobic core collapse, we decide to extract relevant water CVs according to the LDA-like criterion in Eq. 14.

We first calculate the coordination number *C*_WAT,PROT,*i*_ between all protein heavy atoms and the water oxygen atoms with the SWF from Eq. 7. We use *r*_0_ = 3 Å, *n* = 6, *m* = 10, *d*_MAX_ = 10 Å, *r*_NL_ = 15 Åand update the neighbour list every 20 steps. Following a filtering strategy analogous to the one presented above, we determine their discriminative power *S*_*i*_ from Eq. 14 and rank them. We take the 7 highest ranked CVs and bias them progressively along the OneOPES replica ladder.

### Computational details

We use the open-source molecular dynamics software GROMACS 2023 [97] patched with PLUMED2 version 2.9.1 [91, 98] to perform all simulations.

### System preparation for Chignolin

The Chignolin miniprotein has been extensively used for benchmarking new methods aiming at studying fast-folding proteins. In analogy, here we use the double mutant CLN025, shown in Fig. 1 (c), which has two tyrosine mutations (G1Y and G10Y) [65, 66]. The protein is obtained mutating the corresponding wild-type (PDB ID: 1UAO [65]).

Our Python script identifies five hard and eleven soft H-bonds significant in the folded state, and two soft H-bonds significant in the unfolded state. The packing is driven by eight side-chain contacts that are all significant in the folded state.

We use the CHARMM22* force field. The simulation box contains 1907 TIP3P water molecules and two sodium ions to achieve charge neutrality. We use periodic boundary conditions and the particle-mesh-Ewald (PME) method to treat Coulomb long-range electrostatic interactions, while a cut-off distance of 1.0 nm is applied for Coulomb and van der Waals short-range interactions. After 100 ns of simulations of both folded and unfolded states, we run the OneOPES MultiCV simulation in the NVT ensemble with a time step of 2 fs, using the V-rescale thermostat at 340K [99]. The unbiased run is 300 *µ*s long, split into three 100 *µ*s long independent trajectories, while the five OPES and OneOPES runs are each 1 *µ*s long. The free energy results that we present are average over the five independent trajectories.

In the OPES simulations, we bias *s*^HB^ and *s*^SC^ with `OPES METAD EXPLORE` with a `BARRIER` of 20 kJ/mol a deposition `PACE` of 50000 steps (100 ps). `SIGMA` values are the standard deviation of each CV that is extracted from the unbiased folded trajectory. For the OneOPES simulations, the main bias that appears on all replicas is analogous to the one of the OPES simulations (`BARRIER` of 20 kJ/mol, `PACE` of 50000 steps, same `SIGMA`). In higher replicas, starting from replica 1, we sample the multithermal ensemble thanks to OPES MultiThermal, with a `PACE` of 1000. The temperature range for each replica starting from 1 to 7 is: [339K, 342K], [338K, 345K], [337K, 350K], [336K, 360K], [335K, 380K], [330K, 400K], [320K, 420K]. In these replicas, we also bias the water coordination around selected protein heavy atoms. In total we have 4 carbon atoms and 3 oxygen or nitrogen atoms. Replica 1 presents an additional bias on the water coordination of the atom with highest discriminative power, replica 2 independently bias 2 of them until replica 7 that has a bias on all of them. This additional `OPES METAD EXPLORE` bias is deposited with a `PACE` of 100000 steps (200 ps) and a `BARRIER` of 3 kJ/mol. Similarly to the main CVs, `SIGMA` corresponds to the standard deviation in the folded state.

### System preparation for TRP-cage

For the TRP-cage system, we focus on the most studied mutant, namely the Tc10b K8A mutant [67]. The protein is described by the DES-Amber SF1.0 force field [38] and is solvated in a cubic box with TIP4PD water that contains three sodium ions and two chlorine ions to match Ref. [38]. After equilibration, the unbiased run and the enhanced sampling simulations are run in the NPT ensemble with a time step of 2 fs. The temperature is set at 320K and controlled by the V-rescale thermostat [99], while pressure is set at 1 bar and controlled by the C-rescale barostat [100] with a compressibility of 4.5 × 10^−5^ bar^−1^. The unbiased run is 200 *µ*s long, while the five OneOPES runs are each 1 *µ*s long.

Our Python script identifies nine hard and twelve soft H-bonds significant in the folded state, and six soft H-bonds significant in the unfolded state. The packing is driven by twenty-six side-chain contacts that are all significant in the folded state.

In the OneOPES simulations, we bias *s*^HB^ and *s*^SC^ as main CVs with `OPES METAD EXPLORE` on all replicas. The `SIGMA` values are the standard deviation of each CV as extracted from the unbiased folded trajectory. The bias is deposited with a `PACE` of 50000 steps (100 ps) and a `BARRIER` of 30 kJ/mol.

Starting from replica 1, we include OPES MultiThermal, with a `PACE` of 1000. The temperature range for each replica starting from 1 to 7 is: [319K, 325K], [317K, 332K], [315K, 345K], [310K, 360K], [305K, 380K], [298K, 400K], [290K, 420K]. We also bias as additional CVs the water coordination of four carbon and three oxygen/nitrogen atoms. This additional `OPES METAD EXPLORE` bias is deposited with a `PACE` of 100000 steps and a `BARRIER` of 3. Similarly to the main CVs, `SIGMA` corresponds to the standard deviation in the folded state.

### Computational analysis

Cluster analysis on the TRP-Cage trajectories is performed using Visual Molecular Dynamics (VMD)’s *measure cluster* routine [101]. An RMSD threshold value of 1.5 Åis selected considering the number of generated cluster families and the similarity of protein conformations within a cluster family. For each basin, we adapt the PLOT NA routine of the “Drug Discovery Tool” (DDT) to estimate the frequency of occurrence of contacts [102] and assess TRP-Cage’s intra-protein interactions. We set a neighbouring cut-off value of 3.0 Å between interacting residues.

## Data availability

### Code and Data availability

The Python script to generate the features and the enhanced sampling simulation input files are available on Github https://github.com/valeriorizzi/FoldingFeatures. It requires PLUMED [91, 98], version 2.8 or later. The enhanced sampling simulations are run with GROMACS. All the input will also be deposited in the PLUMED NEST repository [103]. The long unbiased reference trajectory will be deposited on Zenodo.

## Acknowledgements

This work was financially supported by the Swiss National Science Foundation and Bridge funding schemes (project numbers: 200021 204795, CRSII5 216587, and 40B2-0 203628). We acknowledge the Swiss National Supercomputing Centre for supercomputer time allocations on Piz Daint and Alps (project ID: s1228, s1274, and lp84). We wish to thank Enrique Alanis Dominguez and Thorben Fröhlking for useful discussions.

