## Supporting Information for "The Arch from the Stones: Understanding Protein Folding Energy Landscapes via Bio-inspired Collective Variables"

### Methods

In the present section, we illustrate the behaviour of the switching function embedded into the CVs. In Fig. S1, we show in blue the switching function from Eq. (3) of the main text for $r_0 = 3\,\text{Å}$, $n = 6$ and $m = 8$, in green the same function for $r_0 = 8\,\text{Å}$, $n = 4$ and $m = 8$, and in red the switching function from Eq. (7) of the main text for $r_0 = 3.5\,\text{Å}$, $n = 2$, $m = 10$, $d_{\text{MAX}} = 5\,\text{Å}$, and $r_{\text{NL}} = 8\,\text{Å}$. In Fig. S2, we focus on the behaviour of the $C'_{HA}$ descriptor for different acceptor-hydrogen distances given a donor-hydrogen-acceptor angle and vice versa, to illustrate the discriminative power of $C'_{HA}$. The reader can refer to Eq. (4-6) and Fig. 1 (a) of the main text for further details.

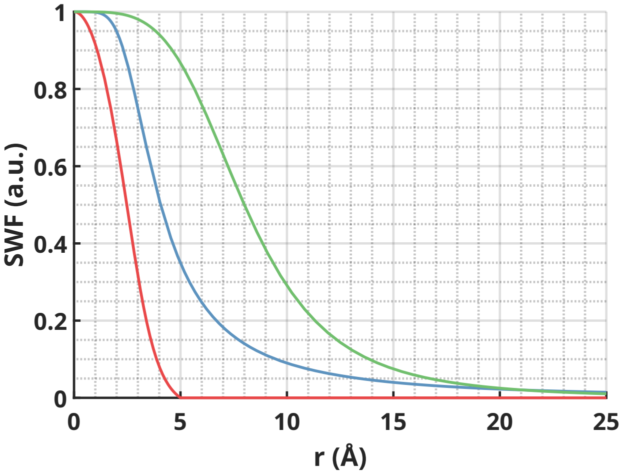


Figure S1: In blue and green, switching function from Eq. (3) of the main text, in red switching function from Eq. (7) of the main text. The corresponding parameters are reported both in the main and in the supplementary text.

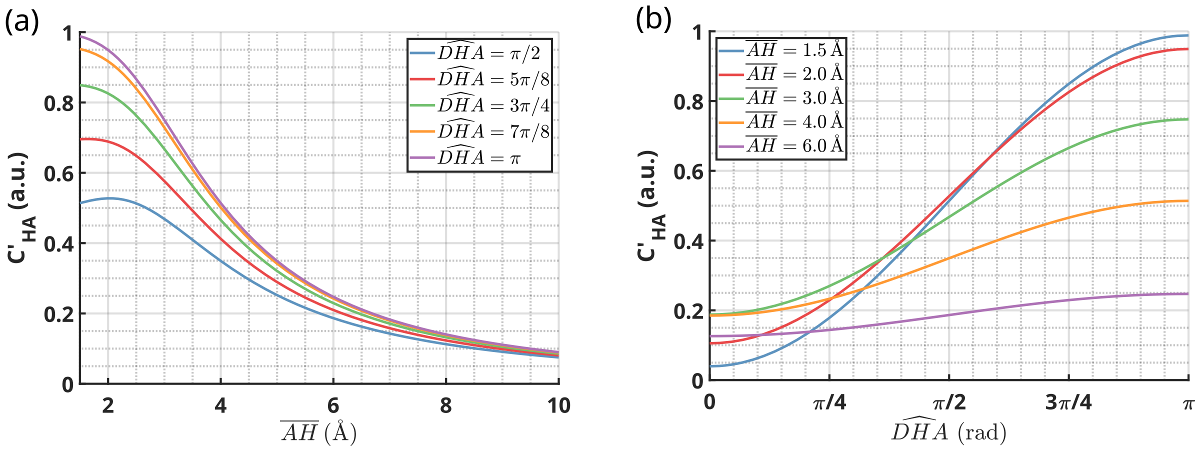


Figure S2: $C'_{HA}$ as defined in Eq. (6) of the main text as a function of (a) the hydrogen-acceptor distance for different donor-hydrogen-acceptor angles and (b) vice versa.

#### Chignolin

In this section, we report additional details of Chignolin's folding process, extracted from the 300 $\mu$s-long unbiased trajectory, the 5 OPES simulations, and the 5 OneOPES simulations. First, we provide further insights into the unbiased simulations, such as the sampling of the RMSD auxiliary CV (see Fig. S3). Then, we performed a thorough assessment of the total energies across the 300 $\mu$s-long unbiased trajectory to ensure the thermodynamic consistency of the simulation (see Fig. S4). Moving on to the biased simulations, we display the exploration of the $s^{HB}$, $s^{SC}$, and RMSD CVs in both the OPES and OneOPES trajectories (see Fig. S5 and Fig. S6, respectively) supporting our claim that the conformational ensemble is thoroughly and consistently explored in both biased and unbiased runs. We conclude this section with a direct comparison of the 2D FES along the $s^{HB}$ and $s^{SC}$ CVs obtained from the OPES and OneOPES enhanced sampling simulations (see Fig. S7).

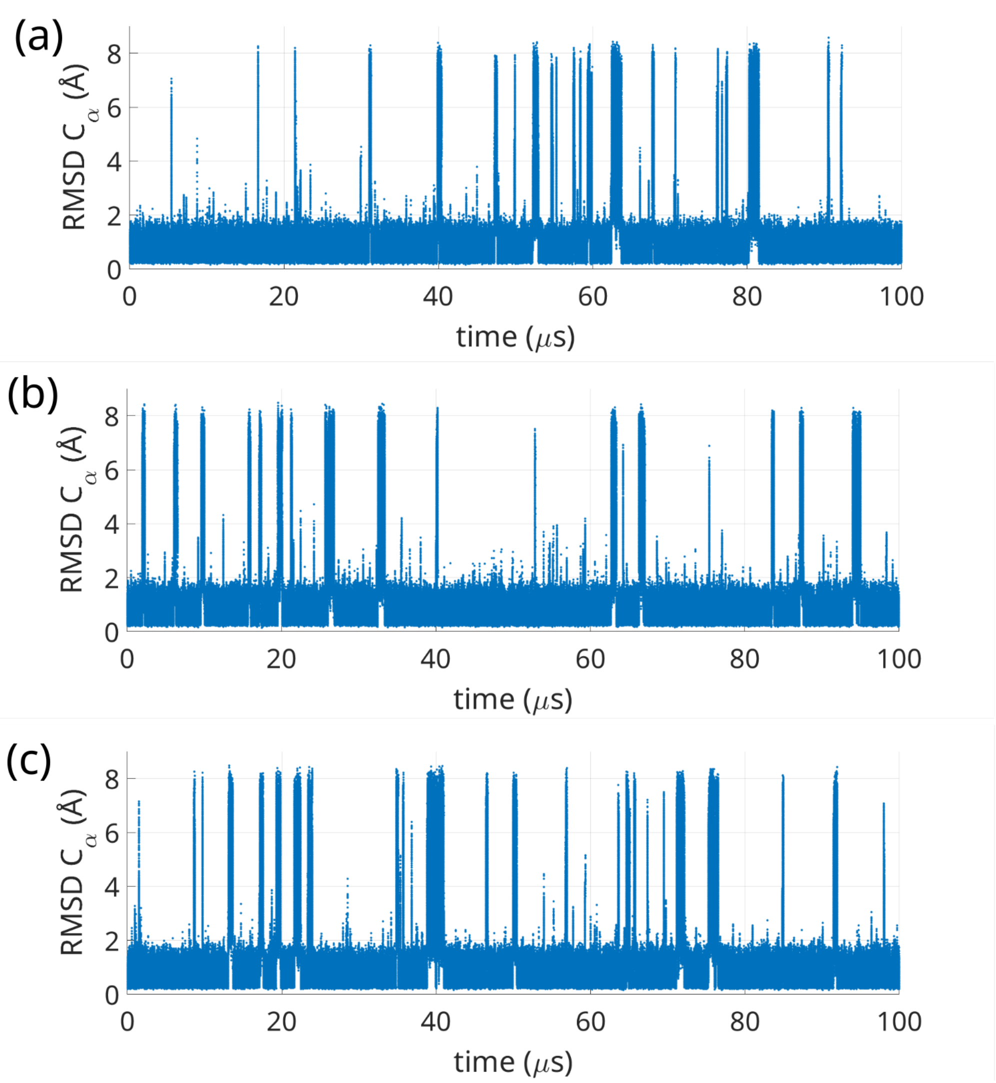


Figure S3: **Folding dynamics of TRP-Cage in unbiased simulations. a-c)** Exploration of the RMSD CV across 300 $\mu$s of unbiased MD simulations. The whole trajectory is made of three independent replicas of 100 $\mu$s each, whose dynamics is displayed in panels (a), (b), and (c).

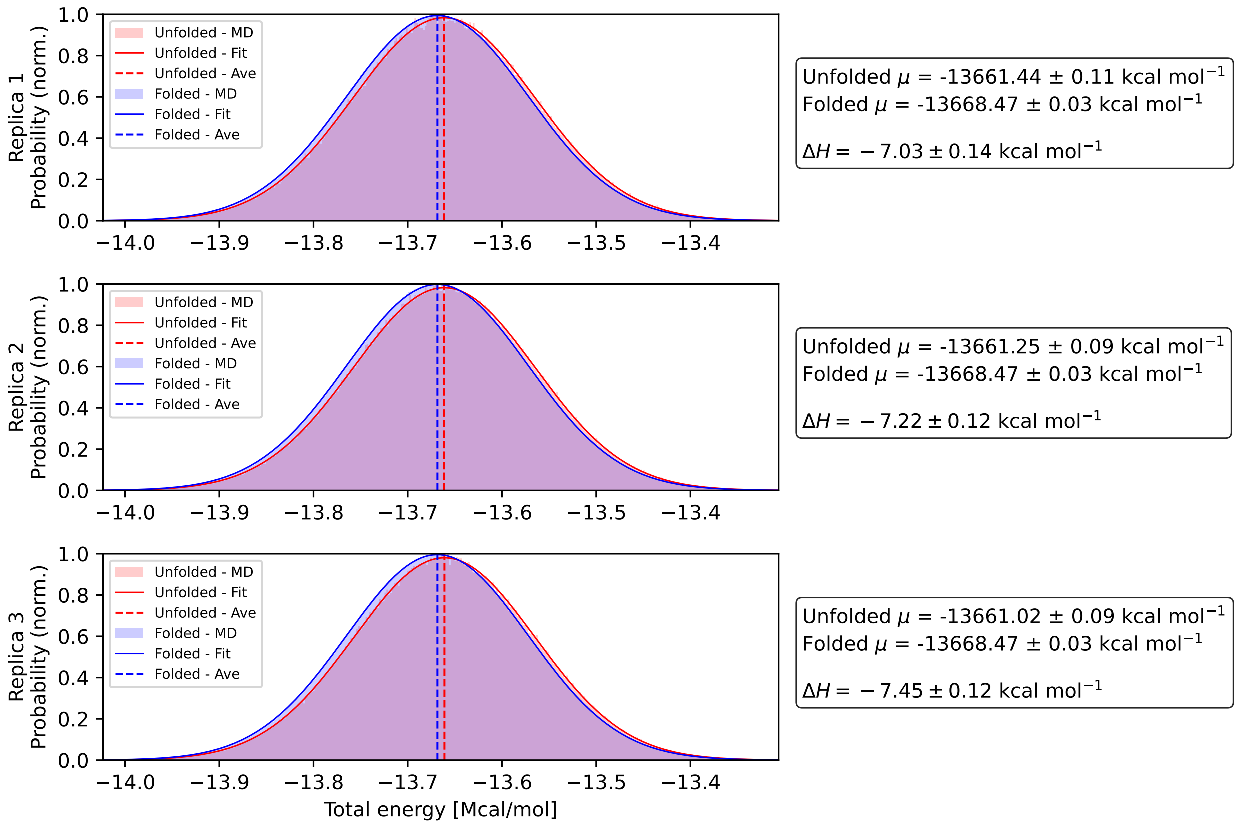


Figure S4: **Distribution of total energies (enthalpies) sampled during the 300 $\mu$s-long unbiased MD simulation of Chignolin.** Histogram representations of enthalpy collected throughout the trajectory, split into the three independent replicas of 100 $\mu$s each. Each distribution is overlaid with a Gaussian fit to highlight the absence of long-timescale drifts or multimodal behaviour.

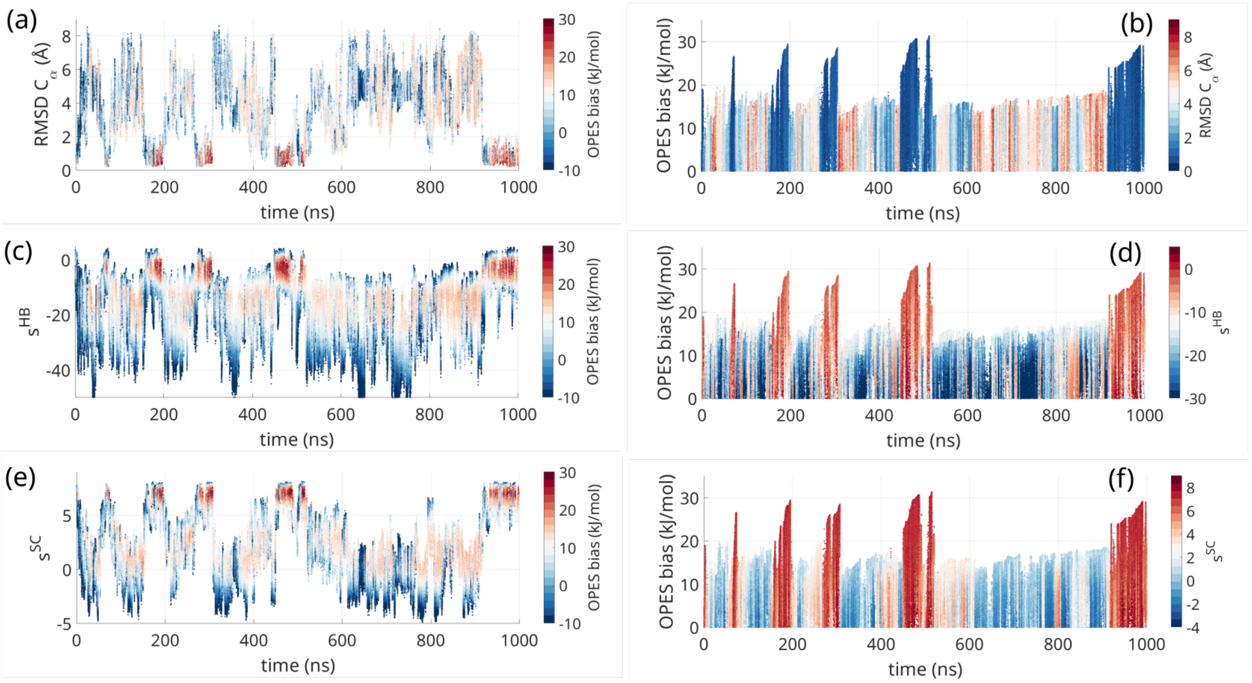


Figure S5: **RMSD and bias evolution during a Chignolin OPES simulation.** **a)** Time evolution of the RMSD CV, coloured by the OPES bias potential. **b)** Time evolution of the OPES bias potential, coloured by RMSD. **c)** Time evolution of the $s^{HB}$ CV, coloured by the OPES bias potential. **d)** Time evolution of the OPES bias potential, coloured by the $s^{HB}$ CV. **e)** Time evolution of the $s^{SC}$ CV, coloured by the OPES bias potential. **f)** Time evolution of the OPES bias potential, coloured by the $s^{SC}$ CV.

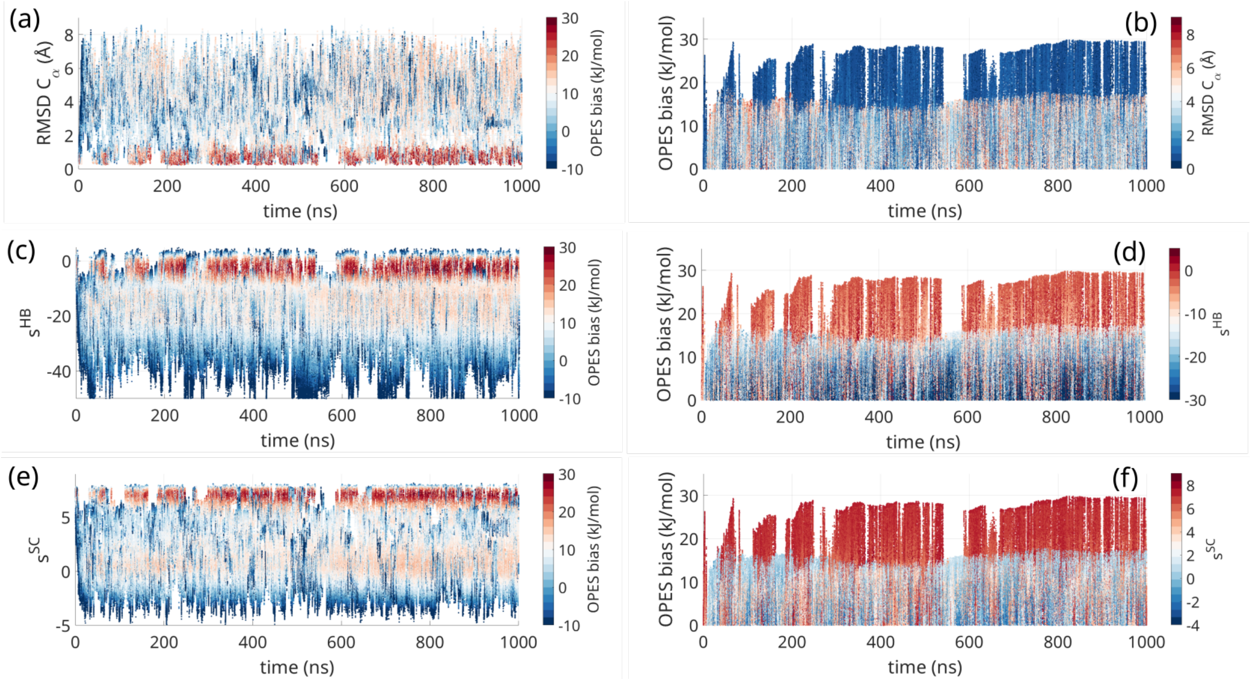


Figure S6: **RMSD and bias evolution during a Chignolin OneOPES simulation.** **a)** Time evolution of the RMSD CV, coloured by the OPES bias potential. **b)** Time evolution of the OPES bias potential, coloured by RMSD. **c)** Time evolution of the $s^{HB}$ CV, coloured by the OPES bias potential. **d)** Time evolution of the OPES bias potential, coloured by the $s^{HB}$ CV. **e)** Time evolution of the $s^{SC}$ CV, coloured by the OPES bias potential. **f)** Time evolution of the OPES bias potential, coloured by the $s^{SC}$ CV.

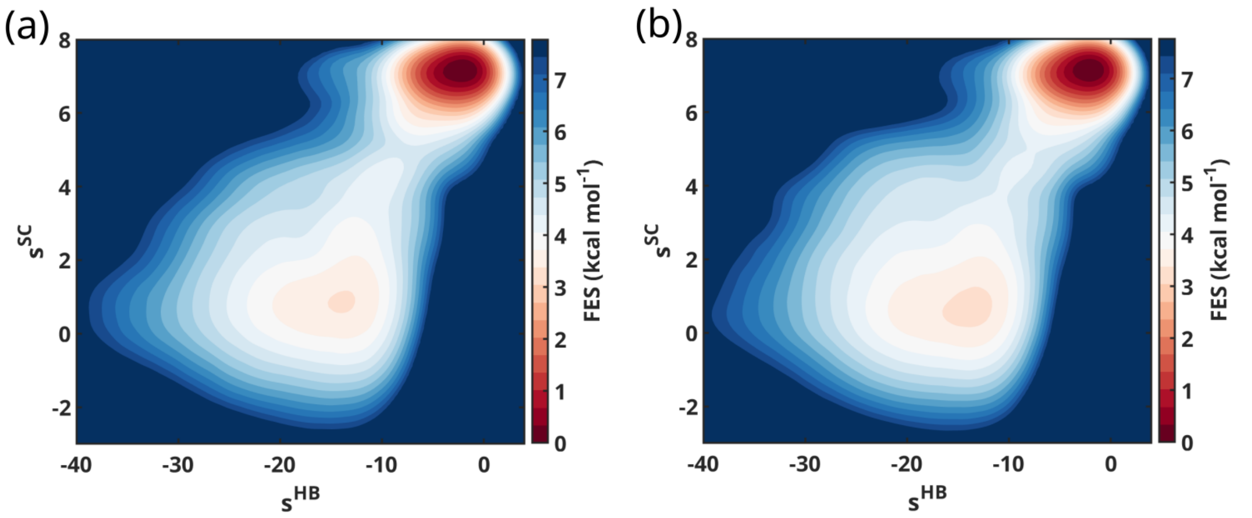


Figure S7: **Chignolin 2D FES as a function of the $s^{HB}$ and $s^{SC}$ CVs.** **a)** Free-energy surface obtained from the OPES simulations. **b)** Free-energy surface obtained from the OneOPES simulations.

#### TRP-Cage

In this section, we report additional details of TRP-Cage folding process, extracted from both the 200 $\mu$s-long unbiased trajectory and the 5 OneOPES simulations. In details, we plan to show additional insights about the sampling of both primary (i.e., $s^{HB}$ and $s^{SC}$) and auxiliary (e.g. RMSD and AlphaRMSD) CVs (see Fig.S8, and Fig.S9), corroborating our assumption that the whole conformational manifold is well sampled in both biased and unbiased runs (see Fig. S10). Lastly, we present further information on the TRP-Cage basins extracted from the 2D FES of the unbiased reference (see Fig. S11) and of OneOPES (see Fig. S12).

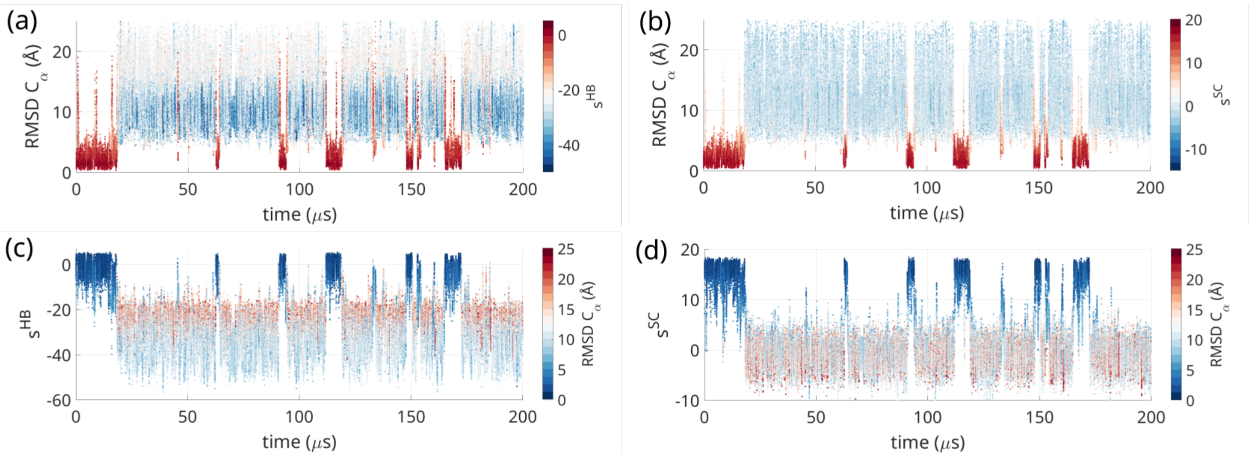


Figure S8: **Folding dynamics of TRP-Cage in an unbiased simulation. a-b)** Time evolution of the RMSD, coloured by the $s^{HB}$ (a) and $s^{SC}$ (b) CVs, respectively. **c)** Time evolution of the $s^{HB}$ CV, coloured by RMSD. **d)** Time evolution of the $s^{SC}$ CV, coloured by RMSD.

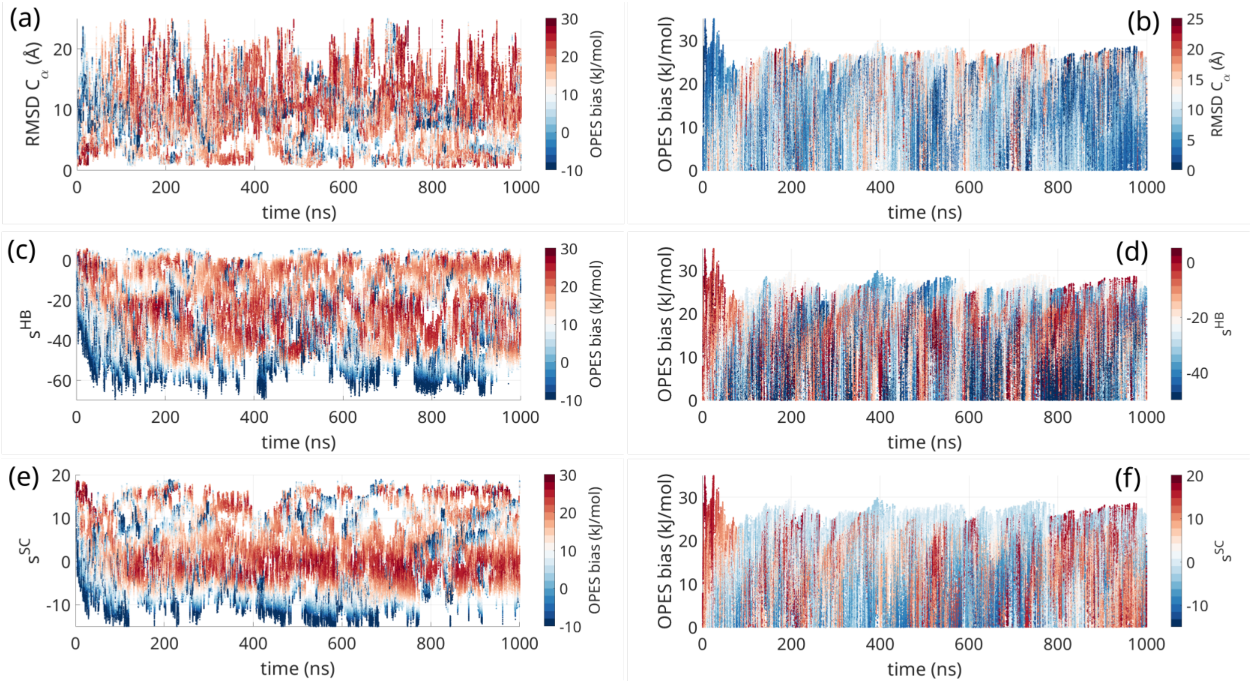


Figure S9: **RMSD and bias evolution of replica 0 during a TRP-cage OneOPES simulation. a)** Time evolution of the RMSD CV, coloured by the OPES bias potential. **b)** Time evolution of the OPES bias potential, coloured by RMSD. **c)** Time evolution of the $s^{HB}$ CV, coloured by the OPES bias potential. **d)** Time evolution of the OPES bias potential, coloured by the $s^{HB}$ CV. **e)** Time evolution of the $s^{SC}$ CV, coloured by the OPES bias potential. **f)** Time evolution of the OPES bias potential, coloured by the $s^{SC}$ CV.

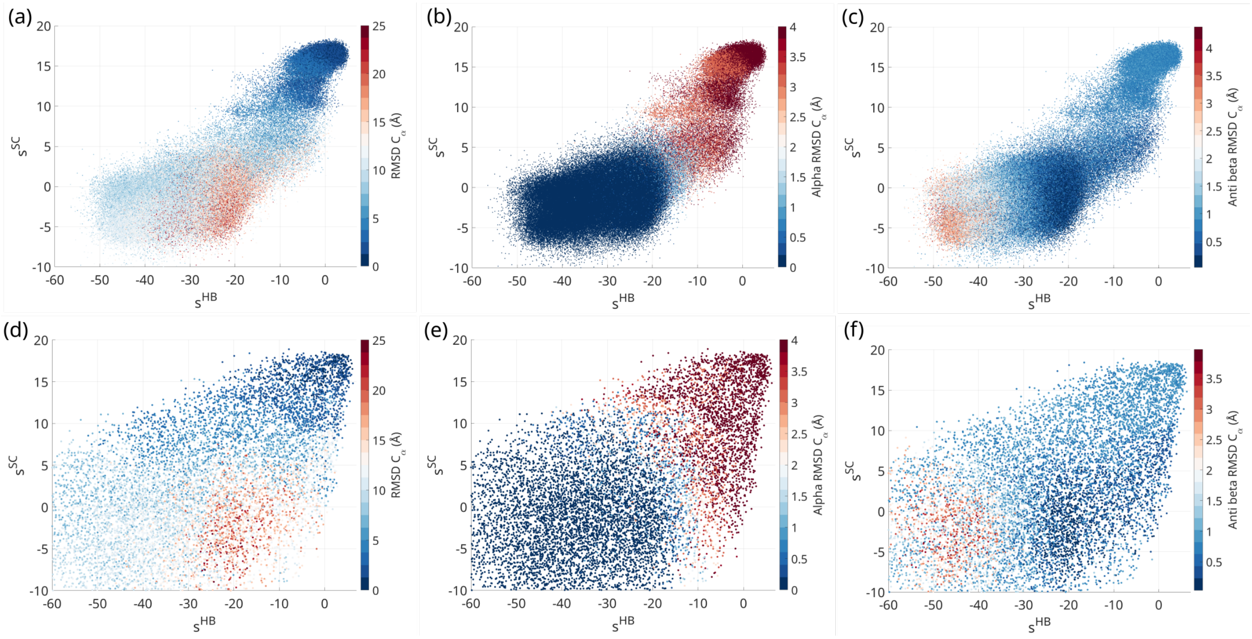


Figure S10: **Comparison of the CV space sampling during unbiased and OneOPES simulations of TRP-Cage. a–c)** Distribution of sampled configurations in the $s^{HB}$–$s^{SC}$ collective variable space during the 200 $\mu$s unbiased MD simulation. **d–f**) Distribution of sampled configurations in the $s^{HB}$–$s^{SC}$ CV space during a OneOPES simulation. For the sake of clarity, points are coloured by different structural descriptors to highlight conformational diversity: panels (a) and (d) are coloured by their RMSD values; panels (b) and (e) are coloured according to their helical content (i.e., AlphaRMSD CV); panels (c) and (f) are coloured according to their $\beta$-structure content (i.e., AntiBeta CV).

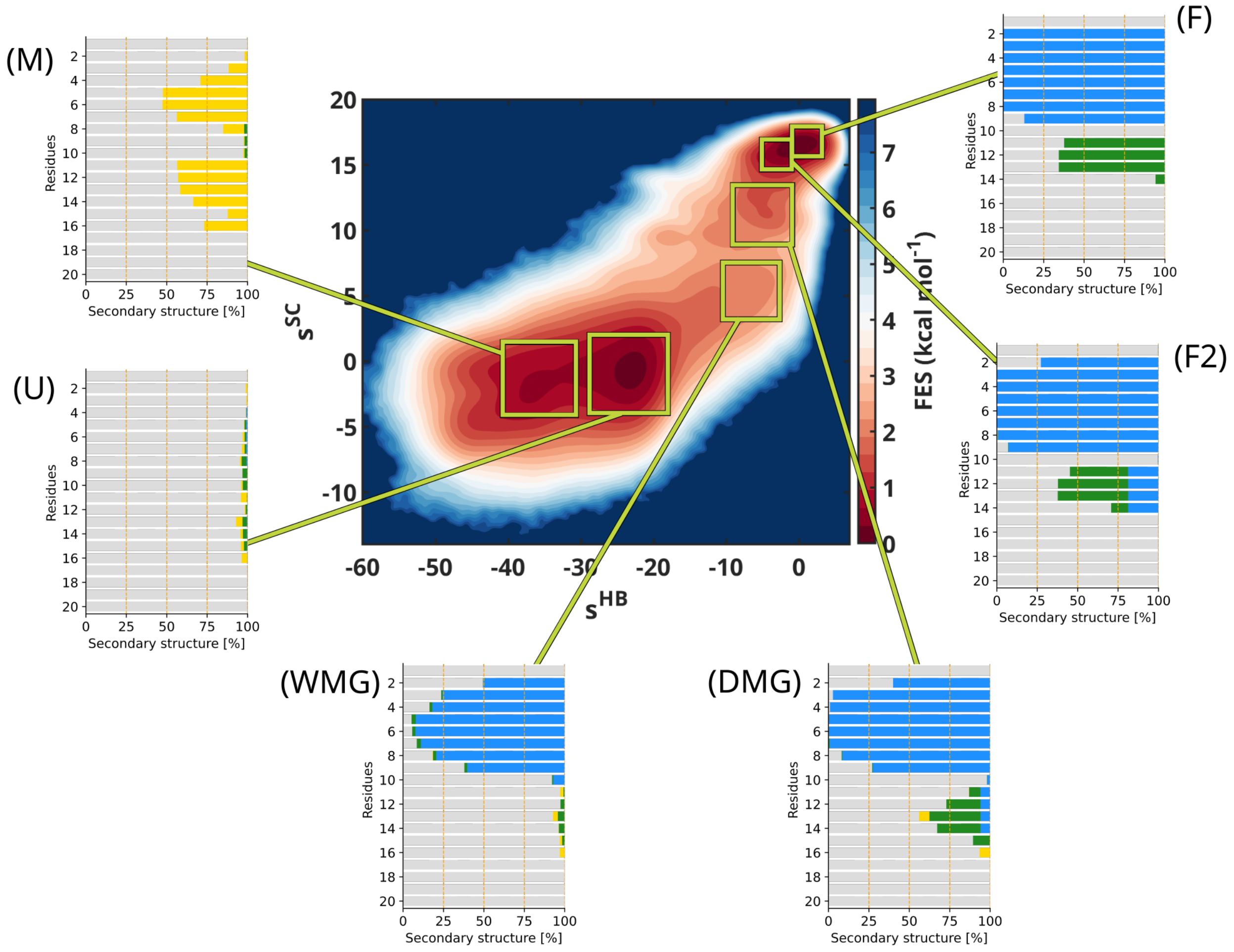


Figure S11: **Trp-Cage folding landscape analysis of the unbiased reference trajectory.** In analogy with Fig. 4 in the main text, we repeat the analysis on the secondary structure of the six conformational ensembles (M, U, WMG, DMG, F2 and F) on our reference unbiased trajectory. For each ensemble, we present a histogram summarising the secondary structure frequency for each amino acid. $\alpha$-helices are coloured in blue, $3_{10}$-helices are coloured in green, $\pi$-helices in magenta, $\beta$-helices in yellow, and random coils in grey.

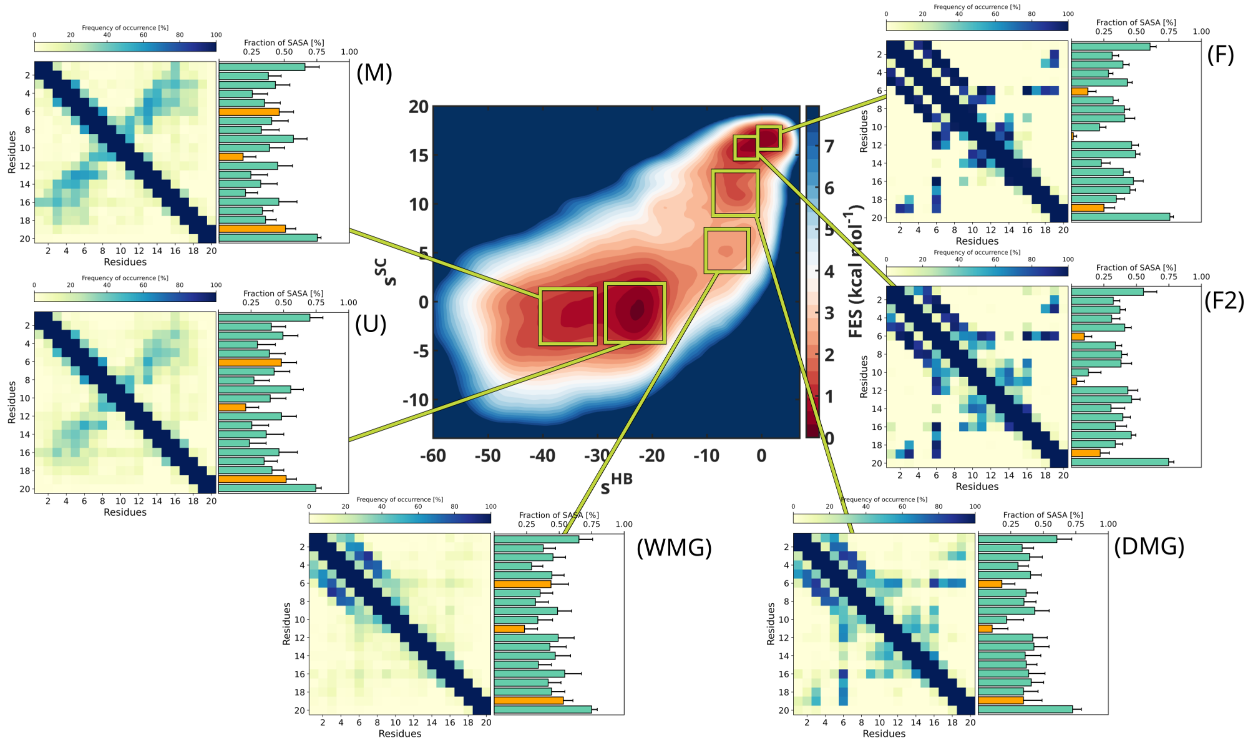


Figure S12: **Additional analysis carried out on the minima extracted from Trp-Cage 2D FES.** For each basin, we display both a frequency map of the inter-residue contacts and the residue-wise histogram of the Solvent-Exposed Surface Area (SASA). In particular, SASA values are normalized with respect to those of the fully solvated amino acids. The contact maps are coloured according to the colour bars over them. In the SASA histograms, bars are shown in green, except for residues W6, G11, and P19, which are highlighted in orange to emphasize their role in hydrophobic core formation
